# Characterization of Z chromosomal inversion and lineage-specific sweeps reveal widespread local adaptation in *Helicoverpa armigera*

**DOI:** 10.1101/2023.09.28.560065

**Authors:** Fushi Ke

**Author notes:** Corresponding author: **Fushi Ke:**.

## Abstract

Efficient pest management requires comprehensive knowledge of its biology, ecology, and evolution, particularly adaptation that exacerbating pest damage. *Helicoverpa armigera* (Hübner) is a notorious pest that attacks more than 200 species, adapts to diverse environments, and has developed resistance to almost all applied insecticides. However, local adaptation of this species was less investigated. Here, we analyzed genomic variation of *Helicoverpa armigera* in China and Oceania to identify adaptative loci in different lineages. Despite intensive gene flow, two differentiated Z chromosomal lineages in Central China (i.e., northern and southern lineages) were formed by chromosome inversion and local adaptation. Based on population genomic analysis, we identified genes related to circadian clock (*period*, *cycle*, *cyr2* and *daywake*), insulin secretion (*IGF2BP2*) and energy metabolism (*SLC25A25*, *CCG7766*, *D11DS*, *TSTP* and *CPQ*) were under selection in northern lineage. These genes may have contributed to its adaptation to high latitudes. In the southern lineage however, the Z chromosome inversion harboring alien alleles of two clock genes (*period* and *cycle*) and may have disrupted clock oscillator for adaptation. We further conducted selective sweep analysis in lineages from Northwestern China, Central China, and Oceania, and found several insecticide resistant genes that shared among different lineages were likely contributed by introgression. Nevertheless, many candidates, including a cadherin that involved in resistance to *Bacillus thuringiensis* protein in Oceanian populations, are lineage-specific. Our results highlight the importance of investigating local adaptation in effective pest control under globalization of agriculture.

## Introduction

Agricultural insect pests cause substantial losses of crop production each year and pose a threat to global food security (Paini et al., 2016). It’s difficult to implement effective pest control strategies without a thorough understanding of pest biology, ecology and evolution, particularly the adaptive evolution that exacerbates the pest damage (Roderick & Navajas, 2003; Renou, 2022). Investigating the ability and genetic basis of pests adapting to local habitats, in terms of adaptation to environments and pesticides thus, would benefit successful pest control. For most insect pests, environmental factors play a crucial role in their performance. Identifying genotypes of higher fitness and selective agents causing environmental adaptation could provide important information for their interaction with local environments, prediction of their further distribution (Pélissié et al., 2018), and implementation of efficient molecular techniques (e.g., through gene editing) for pest control. In the case of resistance to applied pesticides and toxins expressed by transgenic crops, local adaptation could imply that control strategies might need to be tuned according to which populations and regions that are targeted in to keep their efficiency (Alyokhin et al., 2015; Pélissié et al., 2018; Szendrei et al., 2012).

For insect species, forecasting seasonal changes and responding to them in an appropriate way are species’ specific requirements to live in high latitudes or altitudes (Hoikkala & Poikela, 2022). Since temperature and photoperiod are generally reliable indicators of seasonality that have shaped life histories of many temperate climate organisms (Fält-Nardmann et al., 2016), efficient adaptations to thermal stress and photoperiod such as diapause ensure survival and succession of local populations during long overwintering periods (Toepfer et al., 2014). Several pathways have been identified to be underlying the genetic and physiological mechanisms of diapause regulation and evolution (Ragland et al., 2019). Associated genes involved in the circadian clock with diapause were uncovered, with examples showing strong connections between seasonal adaptations and adaptative alleles of clock genes (Ragland et al., 2019; Kozak et al., 2019; Hasebe & Shiga, 2022; Peng et al., 2023). In addition, pathways that insulin involved (Williams et al. 2006; Paaby et al., 2010) and energy metabolism related could also facilitate adaptation of insects to high latitude habitats (Hahn & Denlinger, 2011). Identification of adaptative genes/alleles in the lineages of high latitude thus could give us a comprehensive knowledge of adaptative evolution (e.g., diapause) that could be employed in successful management of insect pests, especially those from temperate regions.

Recombination of genotypes from different genetic background would generate new genetic combinations that facilitate local adaptation of descendants (Ortiz-Barrientos et al., 2016). It could also increase interchange of maladaptative alleles that restrict adaptation of local lineages (Crispo et al., 2011; Arnold et al., 2017). Structural variations such as chromosome inversion that inhibit recombination would restrict gene reshuffling in heterozygotes and maintain adaptative or maladaptative genotypes with linkage (Kirkpatrick & Barton 2006; Roesti et al., 2022). Meanwhile, biotic and abiotic stress that mediated local adaptation would be barrier of gene flow in adaptative loci during homogenization of global genetic variation in differentiated lineages (Tigano et al., 2016). Therefore, investigating the interplay of gene flow, local adaptation and chromosome structural variation would benefit identification of adaptative genes in local populations.

The cotton bollworm, *Helicoverpa armigera* Hübner (Lepidoptera: Noctuidae), is a worldwide crop pest that attacks over 200 species (Pratissoli et al., 2015; Riaz et al., 2021) and shows its capability of adaptation to diverse habitats and hosts. It has also developed resistance to many applied insecticides with a high number of reported cases of insecticide resistance worldwide (Klai et al., 2020). Meanwhile, it could migrate over 2,000 km and mediated intensive gene flow among different populations (Nibouche et al., 1998; Jones et al., 2015; Riaz et al., 2021). Recently, this pest has invaded to South America, hybridized with its sibling species and introduced insecticide-resistant genes into local colonies (Valencia-Montoya et al., 2020). Diapause is an important strategy by which insects, such as *H. armigera*, avoid unfavorable environmental conditions (Danks, 2000). Successful range expansion, particularly at high latitudes, thus requires that species should be able to coordinate their phenology and synchronize life cycles using local temperature and/or day length as cues (Fält-Nardmann et al., 2016; Saikkonen et al., 2012). Investigating phenological adaptation of source populations to environmental cues would help in predicting sites of their invasion and implementing effective pest control. While difference in response to temperature and day length in *H. armigera* lineages have been explored based on molecular experiments (Wu et al., 1997; Wu & Guo 1997; Wu & Guo 2007; Chen et al., 2013; Chen et al., 2016; Virachack et al., 2018), less analysis was conducted to investigate the genetic basis of its acclimation (but see Jin et al., 2023) in the presence of widespread gene flow among geographically distinct lineages.

In this study, we revisited resequencing data of *H. armigera* samples from Asia and Oceania (Zhang et al., 2022), and investigated adaptative genes that have contributed to local adaptation of different lineages. When constructing phylogeny based on Z chromosome polymorphism, we found two differentiated lineages of *H. armigera* in Central China (i.e., the southern (CC-S) and northern (CC-N) lineages) despite widespread gene flow across this region. We further verified intensive introgression between *H. a. conferta* in Oceania and *H. armigera* in Central China, while gene flow of adaptative loci into CC-N was restricted due to local adaptation, representing a model of divergent selection with gene flow (Pinho & Hey, 2010). In addition, our results support Z chromosome inversion harboring *H. a. conferta* alleles of two linked clock genes may have disrupted acclimation in CC-S, and contributed to differential gene flow of *H. a. conferta* into northern and southern *H. armigera* lineages in Central China. Finally, we identify several lineage-specific signals of selection that may be related to insecticide resistance of local populations. We further discuss our findings under current global climate change scenario and severe pest damage to agricultural industry.

## Materials & Methods

### Polymorphic sites and genome resequencing data

We downloaded polymorphic SNPs from figshare (https://figshare.com/articles/dataset/population_genomics_of_cotton_bollworm/17008922). This dataset includes 141 *Helicoverpa armigera* individuals from China, 19 individual *H. a. conferta* from Australia, and nine *H. zea* from North and South America. Samples of China collected from sites at Yellow River Region (YRR), Changjiang River Region (CRR), and Northwestern Region (NR) during 2016-2018. Genomic analysis shows high similarity between samples from YRR and CRR (Zhang et al., 2022), we thus reassigned their geographical origin as Central China (CC). A total of 100 individuals and 41 individuals from CC and NR regions were employed, respectively. Among them, 47, 33 and 10 male individuals with ZZ genotype of sex chromosome, were found in CC, NR *H. armigera* lineages, and *H. a. conferta*, respectively. See Zhang et al., (2022) for detailed information of these samples.

To further calculate population genetic parameters, such as *d*_xy_, we downloaded sequencing reads of 141 *H. armigera* individuals (BioProject No. PRJNA731848) and six *H. zea* individuals NCBI (SRR15526515-20), and called SNPs by using Genome Analysis Toolkit 3.8 (GATK) (DePristo et al. 2011). Sequencing reads of each individual were filtered by using fastq (https://github.com/OpenGene/fastp) and mapped to HaSCD2 assembly (Zhang et al., 2022) using BWA (Li, 2013). The standard GATK pipeline (DePristo et al. 2011) was employed to generate raw sites from bam files. We further conducted filterings in GATK with default settings to exclude low-quality sites, and removed loci with mean coverage depth lower than 3 or higher than 30, or with missing rate higher than 0.5 using VCFtools (Danecek et al. 2011). Unless stated, this dataset with high-quality non-variants was only used for π and *d*_xy_ calculation.

### Phylogeny tree and genetic structure

Phylogeny trees were reconstructed using three methods: neighbour-joining (NJ) method, maximum likelihood (ML) method based on concatenated SNPs, and species tree inferred from windowed trees based on ASTRAL-III (Zhang et al., 2018). For NJ phylogeny, genetic distance matrix of each individual was calculated using VCF2Dis (https://github.com/hewm2008/VCF2Dis) and used for phylogeny construction in phylip-3.697 (http://evolution.genetics.washington.edu/phylip). We further reconstructed the phylogeny relationship based on maximum likelihood (ML) method implemented in IQ-TREE-2.1.3 (Minh et al. 2020). Bi-allelic SNPs thinned by an interval of 100bp were concatenated and used in ML tree construction. We employed +ASC model in model testing to include ascertainment bias correction as SNP data do not contain constant sites. Following the manual, we also appended-bnni to the regular UFBoot command to reduce the risk of overestimating branch support (Hoang et al. 2018). The topologies inferred by NJ and ML methods based on autosomal polymorphism were similar, while topology difference was found when using polymorphic data of Z chromosome. To be specific, we found two differentiated lineages (CC-S, southern lineage, and CC-N, northern lineage) in Central China based on ML tree. Thus, we further constructed the individual phylogeny based on ASTRAL-III using Z chromosomal SNPs. We first divided Z chromosome into 100-kb windows according to coordinates and constructed window tree with SNPs of each window using IQTREE (Minh et al. 2020). A summary tree was then generated based on the constructed 179 window trees in ASTRAL-III (Zhang et al., 2018). Five regions in Z chromosome that were divided by Fst values were further used for tree construction in IQ-TREE.

We further investigated population genetic structure of *H. armigera* (including 19 *H. a. conferta* individuals) based on autosomal SNPs. We generated SNP datasets by pruning SNPs based on LD value of 0.2 with an interval of 50 loci and included only SNP loci at a missing rate ≤ 20% within the intergenic regions 20kb away from the genic regions. We used ADMIXTURE for genetic structure analyses and performed each K from 1 to 20 with 20 replicates. The best K value was identified according to cross-entropy criterion. Employing the same dataset, we conducted Principal Component Analysis (PCA) based on PLINK2 (Chang et al., 2015).

### Inversion identification and genome-wide association analysis

We used Asaph (Nowling et al., 2022) to detect chromosomal inversion in samples collected from Central China as we found two differentiated lineages in phylogeny based on Z chromosomal polymorphims. Asaph could efficiently detect inversion when samples are all from a single population and SNPs are all from a single chromosome (Nowling et al., 2022). Two first PC coordinates of Z chromosomal polymorphism were generated to separate samples in Central China. Further, single-SNP association tests using genotypes of each SNP against the samples’ first PC coordinate were performed. Without long sequencing reads to identify the accurate boundary of the inversion, we set an arbitrary boundary of the inversion by picking SNPs with a-log_10_(P) > 15. Within the inversion, haplotypes possessing derived alleles in over half of the differentiated SNPs, we treated them as haplogroups of inversion. To further identify loci that may have contributed to differentiation of two lineages in China, the GWAS analysis was extended to polymorphic SNPs in autosomal SNPs of each chromosome. We set an arbitrary while widely used *P* value (5 × 10^-8^) as the threshold, and selected SNPs with a *P* value lower than the threshold as outliers.

### Calculation of population genetic parameters

We calculated nucleotide diversity (π), genetic differentiation (*F*_ST_) and genetic divergence (*d*_xy_) by using Simon Martin’s script (https://github.com/simonhmartin/genomics_general). In addition, we calculated window-based parameter *f*_dM_ and *D* based on Dsuite (Malinsky et al., 2021) to investigate genome-wide introgression between different lineages. Only male individuals were included for analysis. Note that we have regenerated a genotype matrix including non-variant sites for accurate calculation of π and *d*_xy_. We further calculated *r*^2^ of SNP pairs in Z chromosome based on PopLDdecay (https://github.com/BGI-shenzhen/PopLDdecay) and compared linkage disequibrilum (LD) between lineages including *H. zea*, CC-N *H. armigera* and CC-S *H. armigera.* To account for different sampling size, we randomly sampled eight individuals in CC-N *H. armigera* and *H. zea* ten times and used these replicates for further comparisons. In addition, to compared LD of different regions, we further divided Z chromosome into “inversion” (identified by Asaph) and “the other” regions, and calculated *r*^2^ of SNP pairs in each region.

### Selective sweep analysis

To identify genetic loci underlying local adaptation in each geographically distinct lineage, selective sweep analysis was performed using site frequency spectrum (SFS) in SWEEPFINDER2 (DeGiorgio et al., 2016). We first assigned *H. zea* as the outgroup and calculated the derived allele frequency spectrum in each population. We further ran selective sweep analysis in each chromosome in a step of 1000 nucleotides. Regions with a CLR value at the top 0.05% of all autosomal regions in each population were identified as outliers. Further, we used a homozygosity-based method and unphased multilocus genotype (MLG) to calculate G12 parameter in each population (Harris et al., 2018). G12 could employ unphased genotype data and is robust similar to H12 (using phased genotype, Garud et al., 2015) in identifying selective sweeps. We don’t intend to separate soft and hard sweeps in each population, but to identify genomic regions under positive natural selection. For *H. armigera* has a similar effective population size while much higher recombination rate (4.86 x 10^-6^ cM/bp, Zhang et al., 2022) compared with *Drosophila melanogaster* (5 x 10^-7^cM/bp, Garud et al., 2015), we used a smaller window of 100 SNPs in a step of 20 SNPs. Windows with G12 values over 99.95% of autosomal windows in each population were identified as outliers.

### Genetic data illustration

For the phylogeny tree, we visualized the tree by using Itol (https://itol.embl.de/). Several R packages, including rworldmap (South, 2011), reshape (Wickham, 2007) and rworldxtra (South, 2012) were used for illustrating inversion and ancestral haplogroups on the map. Pearson correlation was conducted in R and plotted using ggpubr (Kassambara & Kassambara, 2020) and ggplot2 (Wickham, 2011). Package CMplot (Yin, 2020) was used for generation of Manhattan plot. In addition, RectChr (https://github.com/BGI-shenzhen/RectChr) were employed for generating genotype matrix plot.

## Results

### Two differentiated lineages identified by Z chromosomal variation

The phylogeny reconstructed using Z chromosomal variation identified two differentiated lineages in Central China, of which all samples in the southern lineage (CC-S) were from sites near Changjiang River Region (lower latitudes, Figure 1A). This lineage (CC-S) clustered with samples from Oceania (*H. a. conferta*) and differentiated with other *H. armigera* samples in China (Figure 1A). This topology was further supported by summary tree generated by ASTRAL-III based on 179 window (100kb) trees using Z chromosomal polymorphisms (Supplementary Figure 1). The genetic pattern however, was differentiated with the neighbour-joint (NJ) phylogeny (Zhang et al., 2022) and genetic structure (Figure 1B; Supplementary Figure 2) based on autosomal SNPs. To further characterize regional variation across Z chromosome, we calculated genetic differentiation between CC-S and CC-N (CC-N vs. AUS for comparison), and divided Z chromosome into five regions. Of these five regions, two major topologies were identified, with topology one consistent with the autosomal phylogeny (Zhang et al., 2022) and topology two clustering CC-S *H. armigera* lineage with *H. a. conferta*. Tree topology in region 4 was consistent with topology two. Topologies in region 1, region 2 and region 5 that show similarity with topology two have individuals from CC-N or NR clustering with *H. a. conferta* individuals. Only region 3 has a tree topology consistent with topology one based on autosomal SNPs (Supplementary Figure 3). In addition, we found high correlation between Fst_(CC-N vs. CC-S)_ and Fst_(CC-N vs. AUS)_ (Figure 1C), which support widespread introgression of *H. a. confera* into *H. armigera*. This was further supported by window-based introgression analysis (Figure 1D; Supplementary Figure 4).

**Figure 1.**
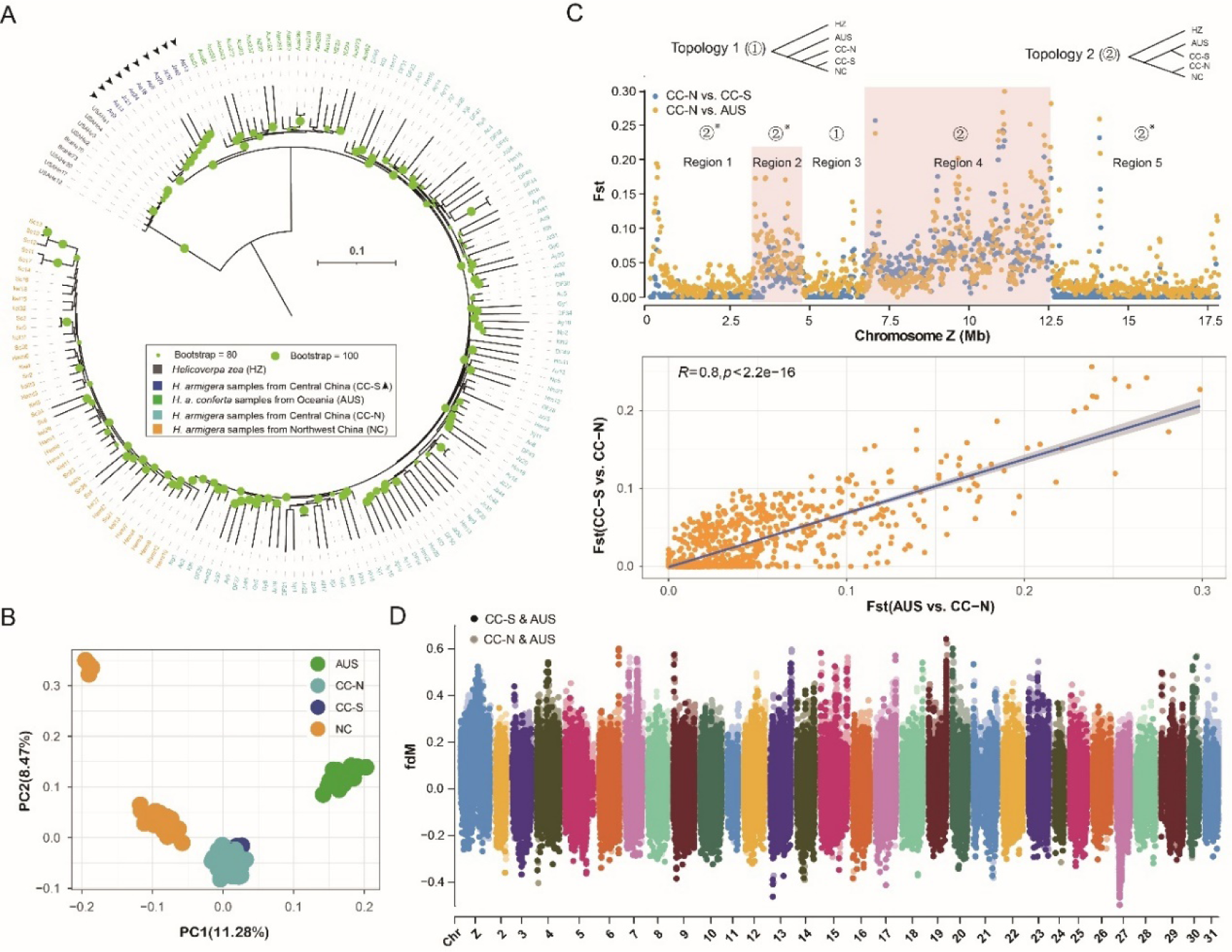
Z chromosome divergence of *H*. *armigera* despite widespread gene flow. (A) Maximum likelihood phylogeny based on concatenated SNPs from Z chromosome. Central China southern lineage (CC-S, with arrows) and Central China northern lineage (CC-N) were labeled with blue and cyan, respectively. Similar topology was reconstructed based on Astral (Supplementary Figure 1) (B) Principal component analysis based on autosomal SNPs. Note the CC-S and CC-N samples were clustered together. (C) Genetic differentiation of *H. armigera* males among different lineages. Two major topologies were illustrated, with topology one (①) consistent with the species tree, while topology two (②) gathering *H. a. conferta* (AUS) and CC-S individuals. “② ※” indicates a window tree similar to topology two, but with CC-S lineage including more individuals from CC-N lineage. See supplementary Figure 2 for more information. The lower panel shows significant correlation between genetic differentiations of CC-S vs. CC-N and AUS vs. CC-N in Z chromosome. (D) Widespread gene flow between *H. a. conferta* from Oceania and *H*. *armigera* populations in Central China.

### Z chromosomal inversion in *H. armigera*

The first principal component (PC) of the Z chromosomal variation divided samples in Central China into two distinguished clusters (Supplementary Figure 5), which is consistent with two differentiated lineages in Central China (Figure 1A). PC1 further identified three individuals (Aq4, Jz42 and DF44) with mixed genetic background. Similar to samples from CC-S, these three individuals were all from the Changjiang River region. Based on the identified PC, we further conducted single-SNP association tests using genotypes each SNP against the samples’ first PC coordinate in Asaph. This analysis identified one distinguished peak at Z chromosome, with the boundary ranges from 6.7 - 12.4 Mb, by setting a threshold of -log10(*P*) > 15 (Figure 2A). We further isolated SNPs with -log_10_(*P*) > 15 in the inversion region and constructed genotype matrix of these differentiated SNPs in male individuals of Central China (Figure 2B, see Supplementary Figure 6 for genotype matrix of all individuals). This inversion region in CC-S lineage was with higher linkage disequilibrilum (LD) compared with other regions of CC-S as well as CC-n lineage and AUS (*H. a. conferta*) (Figure 2C). The genotype matrix identified nine individuals with derived alleles and were with Z chromosomal inversion. The geographical distribution of this inversion showed all individuals possessing this inversion are restricted to Changjiang River region (lower latitudes, Figure 2D).

**Figure 2.**
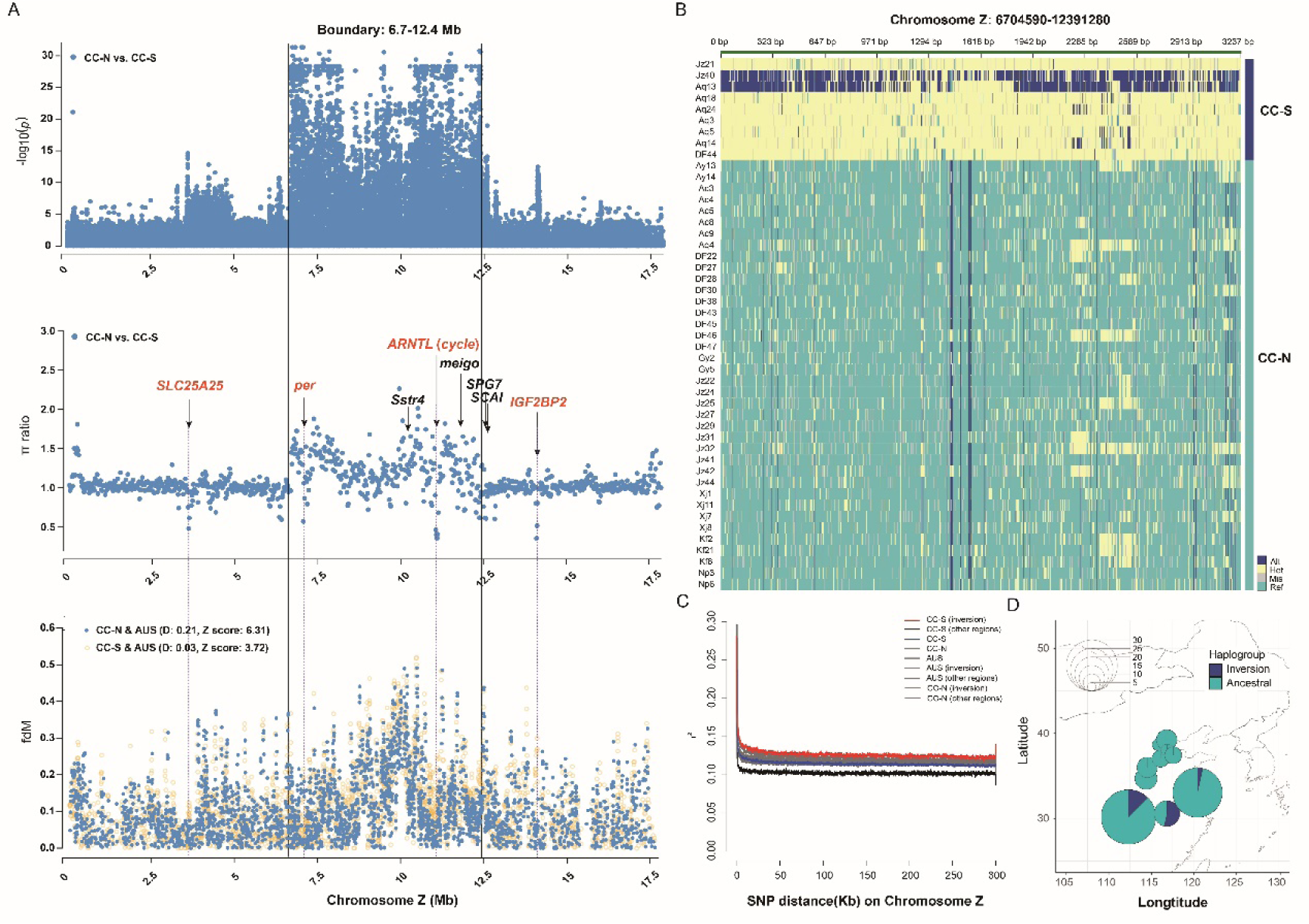
Detection and characterization of putative Z chromosome inversion in *Helicoverpa armigera*. (A) Manhattan plot of GWAS analysis between Z chromosome polymorphism and principal component 1 (PC1). The break point of the inversion was set at the site with -log_10_(*p*) value more than 25. The central panel is π ratio (π_CC-N_/π_CC-S_) calculated in 5kb window, with labeled genes identified in 1% lowest π ratio regions. The lower panel is *fdM* distribution of CC-N vs. AUS and CC-S vs. AUS calculated with 100-SNP window and 25-SNP step. (B) Genotype matrix of highly differentiated SNPs (-log_10_(*p*) > 15) in the putative chromosome inversion. A total of 3,237 SNPs were used for analysis. (C) Linkage disequilibrium analysis in different population and genomic regions; Inversion indicates the putative inversion region, and other regions are the rest of Z chromosome excluding the inversion. AUS are male individuals sampled from *H. a. conferta*. (D) Geographical distribution of two haplogroups (ancestral and inversion) in Central China. Only CC-S and CC-N individuals included for analysis as inversion identified in Central China (see Supplementary Figure 4 for the genotype matrix of all individuals).

Genetic variation of CC-N was higher than CC-S, the regions with lower π ratio (π_CC-N_/π_CC-S_) and higher -log_10_(*P*) thus would be indicatives of selection signals on CC-N lineage (i.e., reduced genetic variation). We treated regions with π ratio lower than 99% of Z chromosome regions (a total of 37 20kb-windows) as candidates and further considered the -log_10_(*P*) more than 7.3 (*P* = 5 x 10^-8^). Interestingly, we identified several genes that maybe potentially important in adaptation of CC-N and differentiation of CC-N and CC-S lineages. Of which, *SLC25A25* and *IGF2BP2* were located outside the inversion, while period circadian protein (*per*) and Aryl hydrocarbon receptor nuclear translocator−like protein 1 (*ARNTL*, or *cycle*) were within the inversion (Figure 2A). Genotype matrix of *per* and *cycle* show that the male individuals in CC-S have similar derived genotype with those in *H. a. conferta* males (Figure 3A). In addition, three male individuals including Aq4, DF44, and Jz22 that are collected from Changjiang River region have the same derived genotypes at both *per* and *cycle*. Several additional individuals from both CC-N (Ay13, DF28, DF46, Xj7, Kf2, Kf21, Kf6) and NR (Hami12, Hami7, Hami9, and Kel26) have only derived genotype in *cycle* (Figure 3A). We also found the genetic differentiation (Fst) and divergence (dxy) of these two genes elevated compared with surrounding regions, and nucleotide diversity of two genes were lower in CC-N lineage compared with those in CC-S lineage (Figure 3B). When conducted protein modeling of two divergent haplotypes dominated in each group, we found difference not only in amino acid but also in protein secondary structure (Figure 3C). In CC-S lineage, these two clock genes, although separated by a distance of around 4 Mb, show highly linkage within and between genes (Figure 3D).

**Figure 3.**
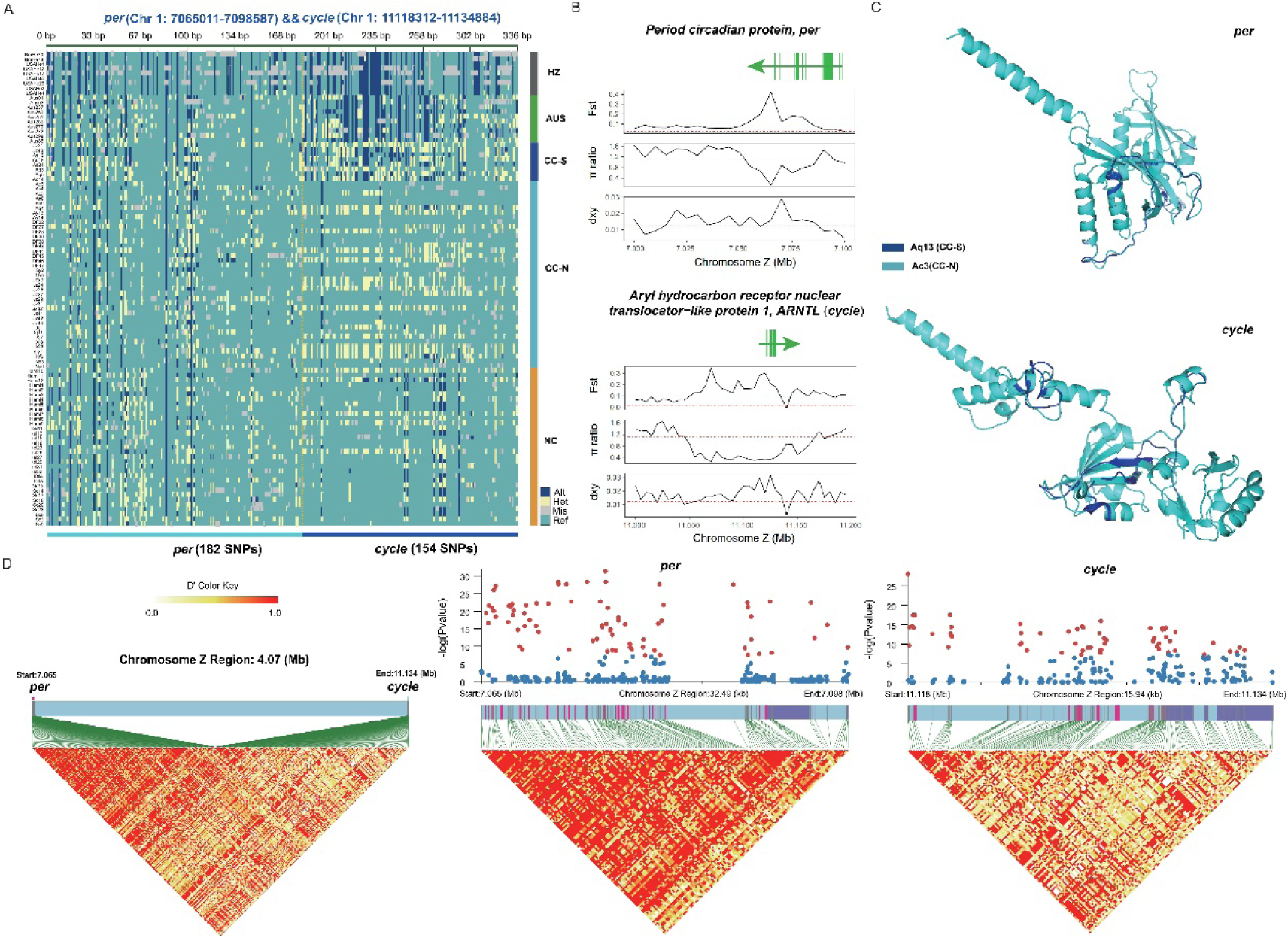
Characterization of *per* and *ARNTL* (*cycle*) in the inversion region. (A) Genotype matrix of polymorphism at CDS regions of *per* and *cycle* genes. (B) Population genetic statistics of *per* and *cycle*. The red dashed line in each panel was the mean value of each statistic calculated by Z chromosomal genetic polymorphism. (C) Protein structures of two differentiated haplotypes in *per* and *cycle*. Cyan structure was constructed from the CC-N dominant haplotype, and additional blue structure that shows the difference was constructed with the CC-N dominant one. (D) Patterns of LD blocks between and within genes (*per* and *cycle*). Red and purple rectangles denote CDS and UTR, respectively. Only male individuals were used in Z chromosome-related analysis.

### Genes contributed to differentiation of southern and northern lineages

In addition to genes in Z chromosome that differentiated between southern and northern *H. armigera* lineages. Genome-wide association analysis of the autosomal SNP genotype against first principal component (of Z chromosome variation) further isolated several regions with significant -log10(*P*) value (> 7.3) and elevated genetic differentiation (Fst). These regions spread across the genome and distributed at more than five chromosomes (Figure 4A). We further checked the annotated genes in these regions, and identified acyl-CoA Delta (11) desaturase (*D11DS*) and bifunctional trehalose-6-phosphate synthase/phosphatase (*TPSP*) in Chr12, phosphorylase b kinase regulatory subunit (*CG7766*), carboxypeptidase Q (*CPQ*) of Chr 13, and cryptochrome-2 (*Cry2*) from Chr15, and circadian clock-controlled protein *daywake* (*annon-3B1.2*) in Chr23. Interestingly, these genes all have elevated Fst and dxy, which supported a divergent selection with gene flow in formation of these divergent peaks. In addition, decreased π ratio (π_CC-N_/π_CC-S_) of these genes compared with nearby regions supported local adaptation of CC-N population may have contributed to this divergence.

**Figure 4.**
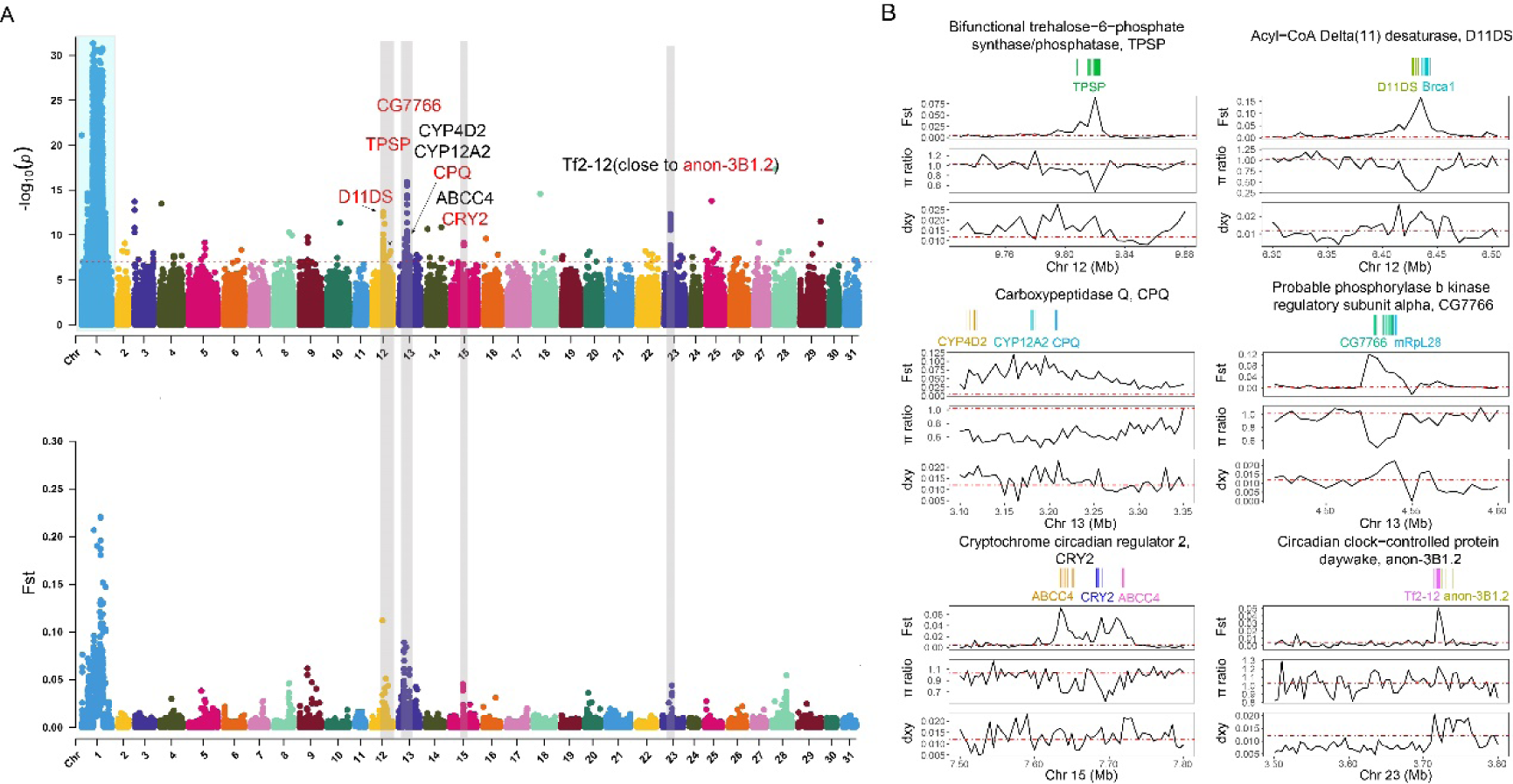
Autosomal genome-wide association tests show differentiation between two *Helicoverpa armigera* lineages. (A) Manhattan plot of genome-wide polymorphisms. Peak regions with more than one highly differentiated SNPs (*p* value lower than 5.0 × 10 ^−8^) were used for gene identification. The lower panel is genetic differentiation (*F*st) between CC-S and CC-N lineages. (B) Characterization of six highly differentiated regions. π ratio indicates π_CC-N_/π_CC-S_, and *d*_xy_ is nucleotide divergence between CC-S and CC-N.

### Widespread selective sweep signals in different *H. armigera* lineages

Several selective sweeps were identified based on strict threshold in different populations (Figure 5). Compared with SWEEPFINDER2, haplotype-based parameter G12 generally identified more regions with sweep signals. Among them, several linked P450 genes identified including CYP337B3V1, CYP6B2, and CYP6a21 were shared across three populations (Figure 5). As we found many regions contributed to differentiation of southern and northern *H. armigera* lineages of Central China, and intensive while diverse application of insecticide for controlling different *H. armigera* populations (Wu et al., 2005; Li et al., 2010; Zhang et al., 2012; Yang et al., 2013; Lu et al., 2021; Downes et al., 2017), different *H. armigera* lineages should have adapted to local climates (Chen et al., 2013) and regionally specific insecticides. Indeed, we found many lineage-specific signals of selective sweeps in populations in Central China, Northwestern China, and Oceania (*H. a. conferta*) based on G12 parameter (Figure 5). In Central China, we found several GSTs on Chr7, carboxylic ester hydrolase (CCE001c) and Juvenile hormone esterase (JHER) of Chr 15, and carboxyl/choline esterase (CCE014a) on Chr30 were under selection. In addition, trehalose-6-phosphate synthase/phosphatase (TPSP) located at Chr12, and Cytochrome P450 12a4 (CYP12A4) and CPQ of Chr13 that were identified as outliers based on G12. In Northwestern China, we found several genes with signals of sweeps except CYP337B3 (Figure 5). In Oceania, cadherin (CadN, Chr 4), choline O-acetyltransferase (ChAT, Chr6), cadherin-89D (Cad89D, Chr6), GST(Chr7), carboxylic ester hydrolase (CCE001c, Chr15), and juvenile hormone esterase (JHER, Chr15) showed selective signals. SWEEPFINDER2 identified less but unique selective signals among different populations. For example, in Central China lineage, PTP10D on Chr13 showed higher CLR values compared with those of other regions. Similarly, two genes (D11DS, and Scd1) on Chr12 have higher CLR values in NC lineage.

**Figure 5.**
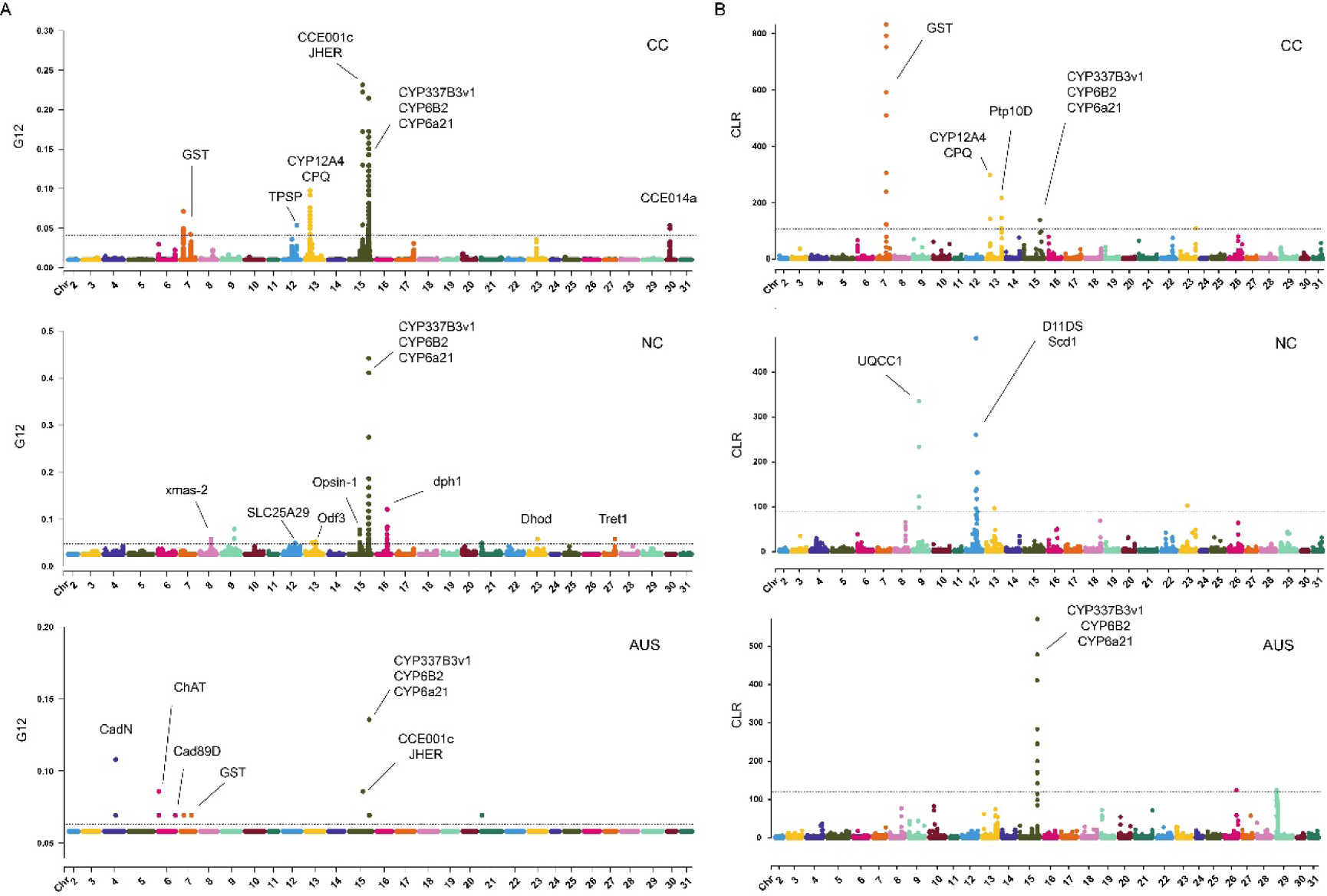
Widespread local adaptation in *Helicoverpa armigera* lineages. Selective sweep signals in different populations from Central China (CC), Northwest China (NC), and Australia (*H. a. conferta*, AUS) based on G12 (A) and CLR (B) statistics. Horizontal dash lines denote threshold values of G12/CLR over 99.95% autosomal windows in each lineage.

## Discussion

Understanding the genetic basis of local adaptation in different insect pest lineages is important in effective pest control and management. To investigate genetic differentiation and local adaptation of two *H. armigera* lineages in Central China identified by Z chromosome polymorphism, we combined genetic differentiation and gene flow analysis to disentangle effects of introgression, local adaptation and chromosome structure on genome-wide genetic variation. Compared with southern lineage in Central China, we identified several genes in northern lineage of Central China with sweep signals (e.g., low nucleotide diversity and high differentiation), restricting introgression from the other subspecies *H*. *a*. *conferta*, and may have contributed to adaptation of this lineage to high latitudes. On the contrary, a Z chromosome inversion in southern lineage, keeping divergent alleles of two circadian clock genes introgressed from *H*. *a*. *conferta*, have contributed to differentiation of these two lineages in Central China, and may have led to disorder of clock oscillator for acclimation. Further, we conducted selective sweep analysis based on two parameters and identified several genes that may have contributed to local adaptation in different *H. armigera* lineages (Northwestern China, Central China, and Oceania). Overall, our results reveal widespread local adaptation of different *H. armigera* lineages that should be considered for predicting *H. armigera* range expansion and executing effective pest management.

### Regional climate contributed to differentiation of southern and northern *H. armigera* lineages in Central China

We conducted analysis on *H. armigera* resequencing data and identify two differentiated lineages in Central China that are differentiated at Z chromosome. Based on population genetic differentiation analysis and gene flow, we found a pattern of divergent selection with gene flow in Central China that was contributed by local adaptation of northern lineage from the high latitudes. A total of four clock genes (Z chromosome: *per* and *cycle*; autosome chromosome: *cry2* and *daywake*) were under selection in the northern lineage. While in the southern lineage, a Z chromosome inversion locked alleles of two core clock genes (*per* and *cycle*) from *H. a. conferta* that further boost divergence of these two lineages. Introduction divergent alleles of these two clock genes into local population may have disrupted sensing system to temperature and photoperiod. In addition, apart from southern lineage, linked introgressed alleles of *per* and *cycle* were only identified in three individuals (Aq4, DF44, and Jz22) from Changjiang River Region without an Z chromosome inversion. These results suggest a combination of certain genotypes in clock genes may be needed to maintain normal function of seasonal sensing system.

Studies shows acclimation to high latitudes through diapause was contributed by multiple loci with minor effects and support a polygenic model (Ragland et al., 2019). In our study, in addition to four clock genes, we further isolated several genes under selection in northern lineage and may have contributed to local adaptation of this lineage to temperature and photoperiod of high latitudes. In Z chromosome, we found *SLC25A25*, a Ca^2+^-sensitive ATP carrier in the inner mitochondrial membrane, fluctuates in a circadian manner in mouse skeletal muscle and might be involve in muscle thermogenesis (Nakao et al., 2017). The other gene on Z chromosome is *IGF2BP2*, which is from insulin-like growth factor2 mRNA binding protein family in human and may associated with impaired insulin secretion (Dai et al., 2020). Phosphorylation of this gene is critical for coordinating cellular function and nutrient metabolism (Dai et al., 2020). In the autosomal GWAS analysis by correlation analysis with each SNP and PCA1 identified by Z chromosome polymorphism, *CCG7766*, a Phosphorylase b kinase regulatory subunit that involved in carbohydrate metabolic process and glycogen metabolic process was identified as an outlier and show local adaptation in northern lineage. The other gene that appears on this gene panel was Acyl-CoA Δ^11^Z-desaturases (D11DS) that occurred ubiquitously in animal and fungal kingdoms and plays essential roles in fatty acid metabolism and response to temperature fluctuations by regulation of cell membrane fluidity (Knipple et al., 1998). In addition, bifunctional trehalose-6-phosphate synthase/phosphatase (TSTP) that shows selective sweep signal in northern lineage is important in energy production and other biological processes of invertebrates (Tang et al., 2018). Additionally, we found an adaptative CPQ gene that encodes carboxypeptidase Q, which is involved in proteolytic functions and resistant to deficit water status (Gheyas et al., 2021; Mohamadipoor Saadatabadi et al., 2021), such as diapause in long overwintering periods.

Overall, our studies identified candidate genes in northern lineage of Central China that involved in circadian clock and could sense seasonal variation of temperature and photoperiod, and in energy metabolism that may have contributed to local adaptation during stressful and long overwintering periods. However, although verified by bioinformatic analysis, additional evidence from transcriptome data and molecular experiment are needed to depict a more robust and comprehensive understanding of adaptation to high latitude of this pest. We also propose a more intensive sampling in southern China as well as southeast Asia to investigate the distribution of Z chromosome inversion and identify genetically distinct lineages that may have adapted to different temperature and photoperiod (Chen et al., 2013; Virachack et al., 2018).

### Adaptation to insecticide leads to widespread local adaptation in different *H. armigera* lineages

Up to now, *H. armigera* shows the highest number of reported cases of insecticide resistance worldwide and have developed resistance against most of applied chemicals and several *Bacillus thuringiensis* (Bt) toxins (Valencia-Montoya et al., 2020). Resistance to insecticides such as carbamates, organophospates, pyrethroids families, or to Bt has been reported all over the world for *H. armigera*, while others including spinosyns, indoxacarb, emamectin, and benzoate were reported regionally (https://irac-online.org/). The differentiation in resistance in regional populations indicate widespread local adaptation to applied insecticides of different *H. armigera* lineages.

*CYP337B3* shows selection signals in all populations, which is consistent with widespread distribution of this P450 gene in field that was resistant to fenvalerate (Joußen et al., 2012; Xu et al., 2014; Han et al., 2015; Rasool et al., 2015). The resistant genotype of *CYP337B3* was also introduced along with populations of *H. armigera* into Brazil (Durigan et al., 2017) and introgressed to its sibling species (i.e., *H. zea*) in South America. In addition, we found selection signals in several lineage-specific genes that indicate local adaptation by development of insecticide resistance. For example, in Central China, the glutathione S-transferase genes that were also identified in previous study (Zhang et al., 2022) took part in insecticide and plant secondary metabolites (Francis et al., 2005; Gawande et al., 2011; Rane et al., 2019). This gene, not identified in Northwestern China (e.g., Xinjiang) but in Oceania, may indicate introgression between *H. armigera* and *H. a. conferta*. In addition, one esterase (*CCE001*) that have involved in fenvalerate resistance was also identified in these two regions (Central China and Oceania), which support widespread introgression of insecticide-resistant genes.

In Central China, one lineage-specific gene encodes carboxyl esterase (*CCE014a*) that could mediate metabolic resistance to organophosphates and carbamates (Venkatesan et al., 2022) and Cytochrome P450 (*CYP12A4*) that could facilitate adaptation to lufenuron (Bogwitz et al., 2005) were identified, while no lineage-specific insecticide resistant genes were isolated in Northwestern China. This may be due to shorter insecticide applied history in Northwestern China compared with that in Central China, differentiation of cropping pattern or geographical and climatic environment (Li et al., 2010; Lu, 2021). On the contrast, we found many population-specific genes (over the threshold) that may have contributed to insecticide resistance in Oceania (i.e., *H. a. conferta* lineage). For example, two cadherins (*CadN* and *Cad89D*) could have contributed to *Bt* resistance (Fabrick & Wu 2015; Du et al., 2019) of transgenic crops in this region were outliers. In addition, one gene that encodes choline acetyltransferase (*ChAT*) could be underlying resistance of this lineage to insecticides that targeted on neurons (Liu et al., 2022). We noted that *Cad89D* and *ChAT* were with weak peaks in Central China (although not being outliers base on threshold), while *CadN* sweep was strong in Oceania and not identified in Central China. The time to start Bt transgeneic crop cultivation in China (since 1997, Huang et al., 2002) and Australia (since 1996, Fitt, 2003) were similar, and the difference in identification of lineage-specific Bt genes could be due to sampling bias or due to the difference in development of *Bt* resistance, which needs further investigation.

### Management of agricultural insect pest required more comprehensive understandings of local adaptation and introgression

For most insect pests, environmental factors are important for successful reproduction and succession of local populations (Fält-Nardmann et al., 2016). Among the genetic agents underlying adaptation of lineages to high latitudes, circadian clock genes could influence insect behaviors such as locomotion, courtship, and photoperiodism should receive more attention (Khyati et al., 2017). In addition, due to more hostile environments of high latitudes, successful range expansion of agricultural pests requires that species are able to use local temperature and/or day length as cues to coordinate their phenology (Saikkonen et al., 2012), pre-adaptation of insects to similar environments thus would facilitate rapid colonization to new habitats (Li et al., 2015). For their importance in viability of insects, techniques such as molecular editing and sterile insect technique (SIT) that involved clock genes (and other genes related to phenological adaptation) should benefit integrated pest management to minimize pesticide use and prevent rapid resistance to insecticides (Khyati et al., 2017).

Addressing the genetic basis of pesticide resistance is the best way to find a sustainable crop plant protection strategy (Barzman et al., 2015). Our analysis of adaptative genes that are related to different insecticides not only identify target genes underlying molecular basis of insecticide resistance, but also revealed that gene flow may have contributed to widespread and rapid development of insecticide resistance of *H. armigera*. This information would help putting forward efficient pest management strategy according to the resistant list of local/regional pest populations. Further analysis to investigate the frequency and distribution of insecticide resistant alleles (introgressed or locally adaptative) across Southeast Asia and Oceania, as well as other regions, are needed for a more successful regional integrated pest management strategy.

### Conclusion

We combined association analysis and population genetic parameters to isolate adaptative agents underlying local adaptation of different *H. armigera* lineages. Genes related acclimation and insecticide resistance were identified despite widespread gene flow. In addition, Z chromosome inversion could disrupt local adaptation by locking maladaptive alleles of introgressed genes. Our results show widespread local adaptation of different *H. armigera* populations, which should further be implemented in integrated pest management and accounted for efficient regional management of insecticide resistance.

---

**Supplementary Figure 1.**
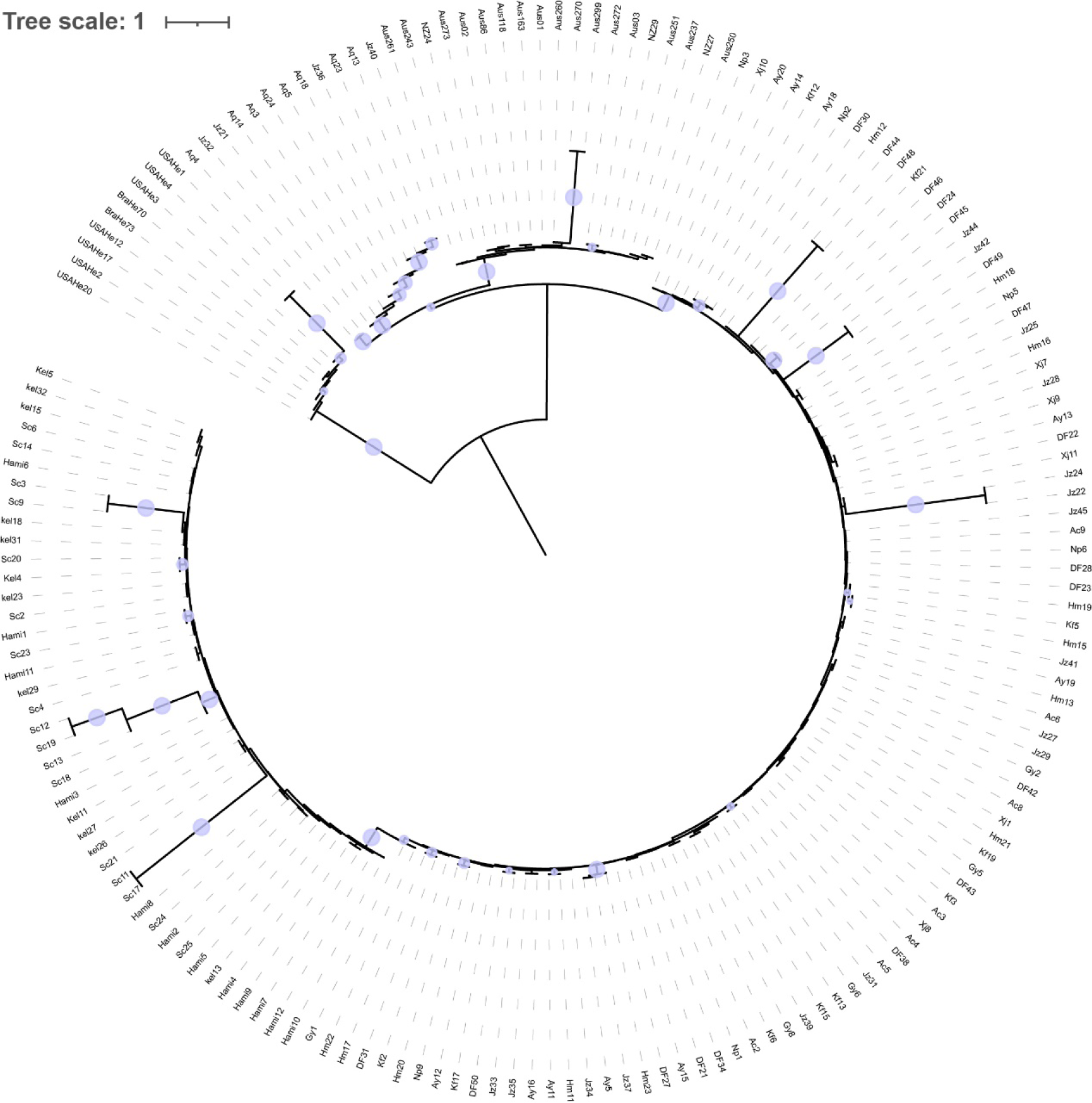
Phylogenetic tree of Z chromosome based on ASTRAL-III.

**Supplementary Figure 2.**
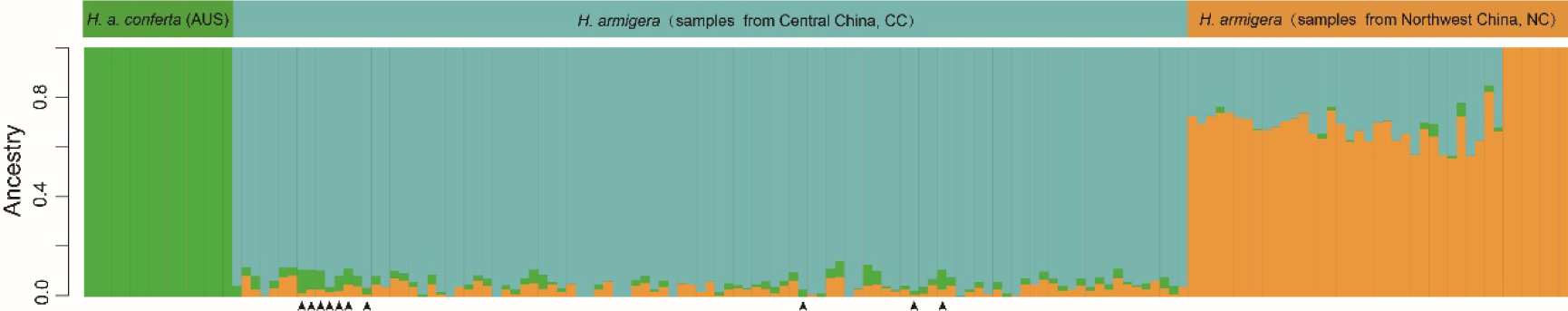
Genetic structure of all *H. armigera* individuals based on autosomal polymorphisms.

**Supplementary Figure 3.**
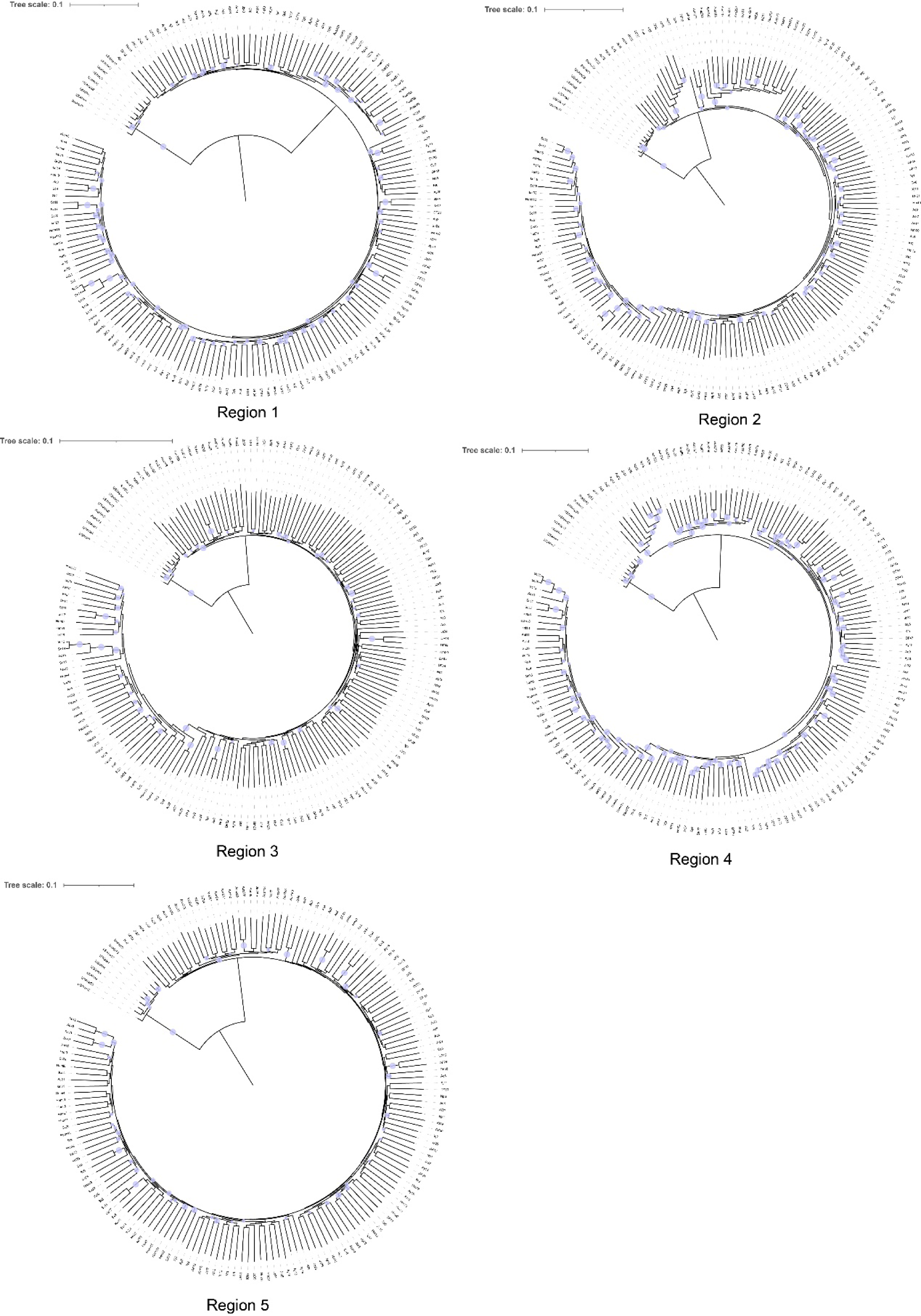
Tree topologies of five regions in Z chromosome.

**Supplementary Figure 4.**
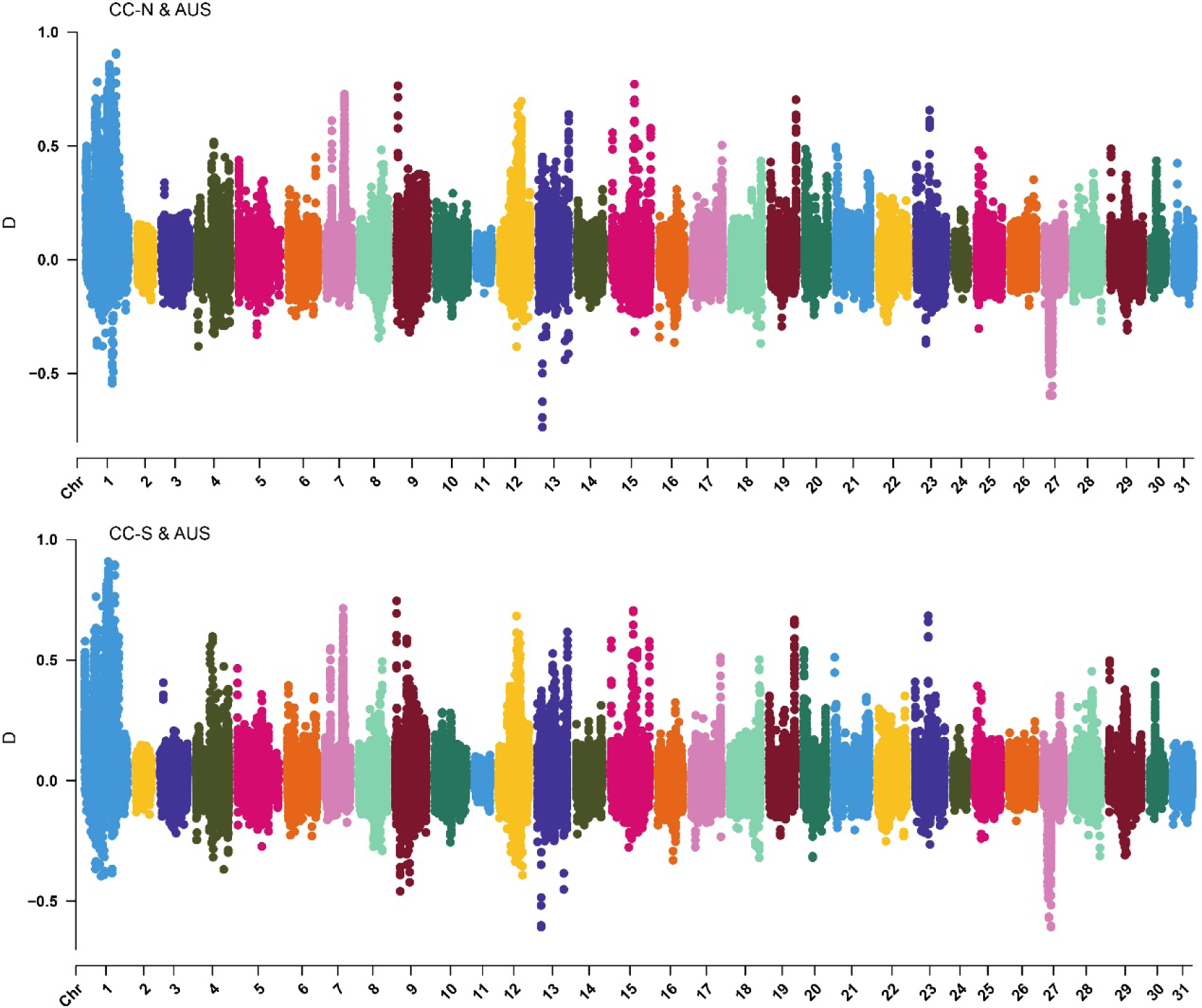
Genome-wide calculation of D value between two *H. armigera* lineages (CC-N and CC-S) in Central China with Oceanian *H. a. conferta* lineage.

**Supplementary Figure 5.**
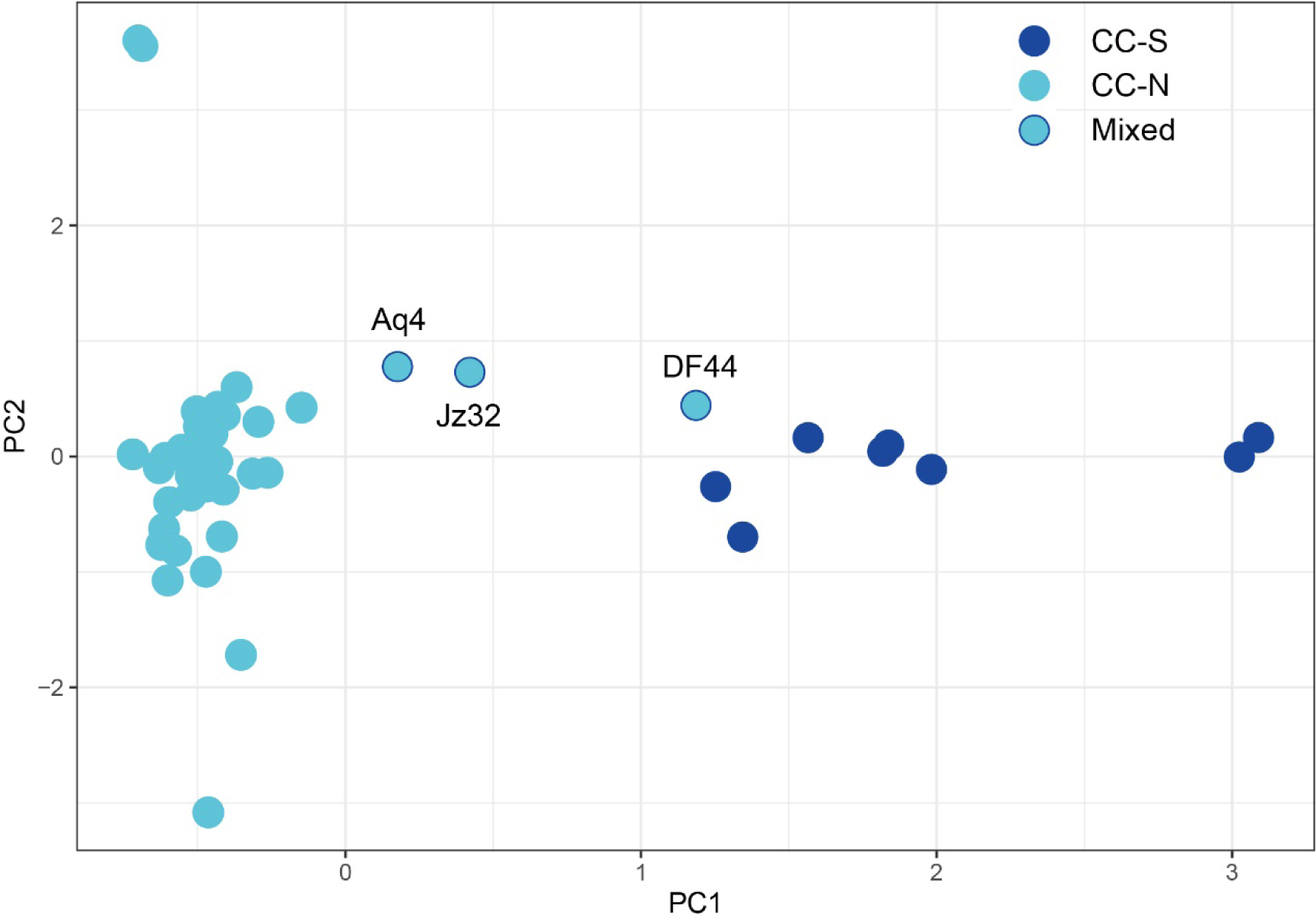
Principal component analysis of *H. armigera* individuals sampled from Central China based on Z chromosomal polymorphisms.

**Supplementary Figure 6.**
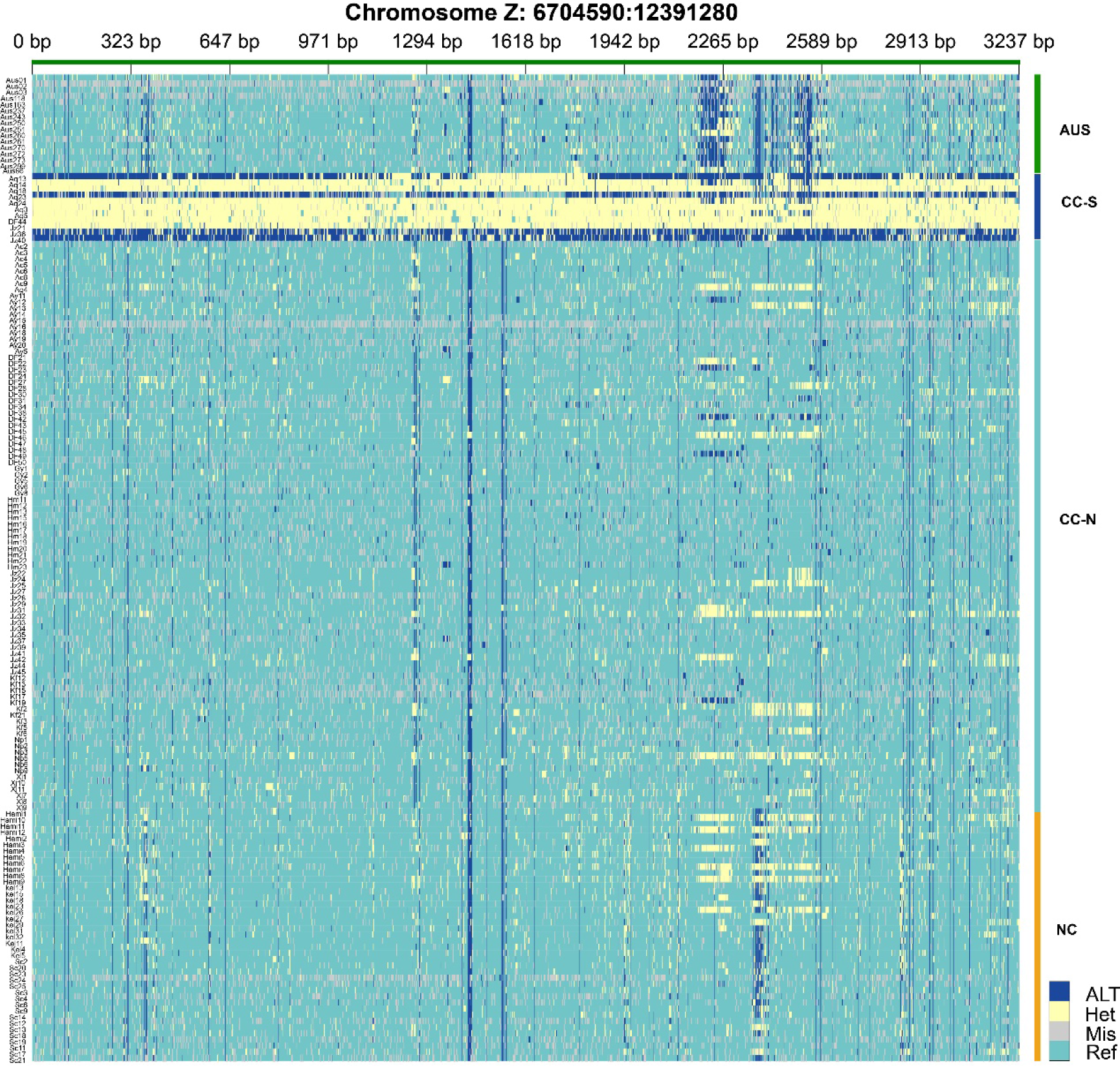
Genotype matrix of all sampled individuals in the putative inversion region of Z chromosome (6.7 - 12.4 Mb).

